# The SARS-CoV-2 Spike mutation D614G increases entry fitness across a range of ACE2 levels, directly outcompetes the wild type, and is preferentially incorporated into trimers

**DOI:** 10.1101/2020.08.25.267500

**Authors:** William A. Michaud, Genevieve M. Boland, S. Alireza Rabi

## Abstract

Early in the current pandemic, the D614G mutation arose in the Spike protein of SARS-CoV-2 and quickly became the dominant variant globally. Mounting evidence suggests D614G enhances viral entry. Here we use a direct competition assay with single-cycle viruses to show that D614G outcompetes the wildtype. We developed a cell line with inducible ACE2 expression to confirm that D614G more efficiently enters cells with ACE2 levels spanning the different primary cells targeted by SARS-CoV-2. Using a new assay for crosslinking and directly extracting Spike trimers from the pseudovirus surface, we found an increase in trimerization efficiency and viral incorporation of D614G protomers. Our findings suggest that D614G increases infection of cells expressing a wide range of ACE2, and informs the mechanism underlying enhanced entry. The tools developed here can be broadly applied to study other Spike variants and SARS-CoV-2 entry, to inform functional studies of viral evolution and vaccine development.

## Introduction

In January of 2019, a new strain of coronavirus, SARS-CoV-2, was identified as the cause of a severe acute respiratory illness in Wuhan, China. Since then, SARS-CoV-2 and the associated disease, COVID-19, have imposed a devastating toll on all corners of the world. A critical aspect of the scientific community’s response to this global pandemic has been tracking the viral variants that emerge worldwide. Towards this aim, an amino acid change in the Spike protein of the SARS-CoV-2 was identified, which was originally found in only a minor percentage of the isolated sequences but by March 31, 2020 was the dominant form globally^1^. The Spike protein of the coronaviruses is synthesized as a polyprotein and is cleaved into two separate subunits, S1 and S2, by cellular serine proteases ^2,3^. Three processed S protomers interact to form a homotrimer on the surface of the virus ^4,5^. This trimer mediates the binding of the viral particles to the cellular receptor, angiotensin-converting enzyme 2 (ACE2), and the subsequent fusion to and entry into the target cells. The new variant, D614G, has an aspartic acid to glycine substitution in the S1 subunit of the Spike protein 66 amino acids upstream of the S1/S2 cleavage site ^1,3^.

While the founder effect - whereby a small number of strains establish a new population - is a possible explanation for the increasing prevalence of the G614 variant ^6^, accumulating evidence points to an evolutionary fitness advantage conferred on SARS-CoV-2 by this mutation. First, sequence analysis has revealed that the D614 variant was well established within several populations before the arrival of the G614 variant ^1^. Second, hospitalized patients infected with the G614 variant likely have a higher viral load at the time of diagnosis compared to the D614 variant ^1^. Finally, in vitro experiments using pseudotyped viruses and viral-like particles have revealed that the G614 variant has a greater entry fitness compared to the D614 variant ^1,7,8–10^. However, several key questions remain unanswered. First, it is unclear if the G614 variant can outcompete the D614 variant in a physiological setting. Second, whereas in the human body SARS-CoV-2 infects cells that differ widely in their level of ACE2 expression ^11–13^, the prior experiments utilized cell lines expressing an artificially high and fixed level of ACE2 to demonstrate entry fitness differences amongst different strains ^1,7,14,15,9,10^. Finally, the mechanism by which the D614G mutation enhances entry into the target cells is unknown. Recent unreviewed data by Zhang et al. ^7^ demonstrated that pseudotyped virions can package more of the G614 Spike protein compared to the D614 variant. Also, it seems that the association between the S1/S2 subunits is stronger in virions containing the G614 variant. Cryoelectron microscopy (Cryo-EM) structures suggest that the side chains of the D614 and T859 residues from neighboring Spike protomer form hydrogen bonds ^1,4,5^. In the G614 variant, this hydrogen bond is not present. Therefore, the D614G mutation appears to fundamentally change the protomeric interactions within a Spike trimer. The consequence of this change on the stoichiometry of the Spike trimers and the subsequent viral entry is unknown.

In this work, we set out to answer these questions. First, to quantify the relative fitness advantage of the G614 over the D614 variant, we set up an in vitro competition assay in which lentiviral vectors expressing different fluorophores and pseudotyped with either S^D614^ or S^G614^ compete for entry into the same cells, mimicking the situation when two different viral strains infect the same host and compete for binding to target cell receptors. Second, to investigate the fitness advantage of the D614G mutation when infecting different cell types with varying expression levels of ACE2, we made an inducible HEK293T cell line expressing the ACE2 receptor under the control of doxycycline. In these cells the ACE2 expression levels is adjustable and can mimic levels encountered in vivo across different tissues and physiological conditions. Finally, we investigated the mechanism of the G614 variant’s enhanced entry fitness compared to the D614 virus. We synthesized lentiviral particles with chimeric Spike trimers containing both the S^D614^ and S^G614^ monomers. Following stabilization of the Spike trimers using a crosslinking reagent, we were able to extract and purify them from the surface of the virions by immunoprecipitation using C-terminal tags fused to the S^D614^ and S^G614^ monomers. This allowed us to analyze the relative amount of each protomer in the chimeric Spike trimers.

Due to the spike protein’s central role in the viral life cycle and its prominence in inducing immunity against the virus and the ongoing vaccine development efforts, it is essential to develop quantitative in vitro techniques to study these variants. In addition to providing evidence for the evolutionary fitness of the D614G variant and elucidating the mechanism underlying its enhanced entry, the tools developed in this work can be applied to the study of the future spike variants and the process of entry by SARS-CoV-2 more broadly.

## Methods

### Cell lines

Clonal HEK293T cells expressing ACE2, HEK-ACE2, are a gift from Michael Farzan via the Massachusetts Consortium on Pathogen Readiness. These cells were made by transducing HEK293T cells with an exogenous ace2 gene. A clone with high expression of ACE2 was selected ^14^.

### Primers

All primers used in the PCR and sequencing reactions are summarized in Table 1 (supplementary material).

### Plasmids and viral constructs

The psPAX2 plasmid was a gift from Didier Trono (Addgene plasmid # 12260; http://n2t.net/addgene:12260; RRID:Addgene_12260) and is a lentiviral packaging vector used for second- and third-generation lentiviral production systems. The pSin-DsRed-IRES-Puro and the corresponding GFP-expressing vector were made using pSin-EF-Sox2-Puro, which was a gift from James Thomson (Addgene plasmid # 16577; http://n2t.net/addgene:16577; RRID:Addgene_16577) ^16^. To make the backbone vector, we removed the sox2 ORF from the original vector using the EcoRI and BamHI restriction enzymes. Using the standard “cut-and-paste” method, we cloned an oligonucleotide (ggaattcgtttaaacggatcc) containing the PmeI cut site flanked by the EcoRI and BamHI sites into the cut pSin vector to make the circular vector, pSin-EPB-IRES-Puro. The dsred and gfp ORFs were PCR amplified from and subcloned into the pSin-EPB-IRES-Puro using the EcoRI and BamHI sites to create the pSin-DsRed-IRES-Puro and pSin-GFP-IRES-Puro vectors, respectively.

The vectors expressing the Spike proteins are a modification of a PiggyBac (PB) vector generously provided by Sahand Hormoz ^17^. Using the CoV-10 and CoV-24 primers, the codon-optimized spike gene was PCR-amplified from a plasmid obtained from Sino Biological (VG40589-UT). This gene encodes a version of the Spike protein with amino acid identical to QHD43416.1 GenBank entry. The resulting gene, S-D614-ΔC-HA, lacks the C-terminal 19 amino acids in the Spike protein and contains a C-terminal HA tag (YPYDVPDYA). We used the NEBuilder^®^ HiFi DNA Assembly Cloning Kit from the New England Biolabs for cloning the S-D614-ΔC-HA amplicon into the PB vector. First, we further PCR-amplified the S-D614-ΔC-HA with the primers CoV-20 and CoV-21. This round of amplification adds the appropriate overhang sequences to the amplicon’s ends for the assembly process. Next, we linearized the parental PB vector using the restriction enzymes NheI and HindIII. The linearized vector and the amplicon with the appropriate overhang were used in the DNA assembly reaction according to the manufacturer’s protocol. The resulting vector was transformed into One Shot^®^ TOP10 chemically competent E. coli, and the resulting colonies were screened for the presence of the S-D614-ΔC-HA sequence using the sequencing primer, CoV-31. This vector, PB-S-D614-ΔC-HA, expresses the with the C-terminal 19 amino acids deleted. An HA tag is inserted in the C-terminus instead.

The PB-S-G614-ΔC-HA vector is obtained from the PB-S-D614-ΔC-HA vector using the Quickchange XL Site-directed Mutagenesis (Agilent) and the primers CoV-35 and CoV-36. The resulting D614G mutation is confirmed by sanger sequencing using the CoV-37 primer.

To add the 3xFLAG tag to the Spike protein instead of the HA tag, we amplified the spike gene from the Sino Biological vector using three subsequent rounds of PCR using primer pairs {CoV-45 and CoV-42}, followed by {CoV-45 and CoV-43}, and finally, {CoV-45 and CoV-46}. The resulting amplicon was digested by the restriction enzymes NheI and HindIII. The PB-S-D614-ΔC-HA was also digested by the same restriction enzymes to drop out the S-D614-ΔC-HA insert. The Flag-tagged spike gene was ligated to the linearized vector using the T4 DNA Ligase enzyme from NEB. The resulting plasmid, PB-S-D614-ΔC-3xFLAG, was confirmed with sanger sequencing.

The inducible ACE2 vector, PB-Tet-hAce2, was made from the XLone-GFP parental plasmid which was a gift from Xiaojun Lian ^18^ (Addgene plasmid # 96930; http://n2t.net/addgene:96930; RRID:Addgene_96930). The ace2 gene was amplified from a plasmid gifted by Hyeryun Choe ^14^ (Addgene plasmid # 1786; http://n2t.net/addgene:1786; RRID:Addgene_1786) using CoV-39 and CoV-40 primers. The amplicon was then digested with the SpeI and KpnI restriction enzymes and ligated into the similarly-digested XLone-GFP empty vector.

The pTransposase which expresses the piggyBac Transposase enzyme which mediates stable integration of the PB vectors into the cellular genome was a gift from Sahand Hormoz ^17^.

### Surface staining of the Spike-producing cells by solubilized ACE2

HEK293T cells were transfected in 6 well plates with 2 μg of PB-S-G614-ΔC-HA and PB-S-D614-ΔC-HA plasmids in triplicates. Forty-eight hours later, the cells were removed from the plate using 5mM EDTA solution and stained with a solubilized ACE2 molecule carrying a human IgG1 Fc at the C-terminus (Acro Biosystems, AC2-H5257). After 45 minutes, the cells were washed twice and incubated with an R-phycoerythrin (PE)- conjugated mouse anti-human IgG1 secondary antibody (BD Biosciences, 555787) on ice for 30 minutes. After washing away the unbound antibodies, the Spike protein expression level was quantified as the mean of the fluorescent intensity (MFI) in the PE channel using flow cytometry (BD Accuri™ C6).

### Pseudotyped virus production and quantification

HEK293T cells were seeded in T150 or T75 flasks at ~50% confluency the night prior to transfection. The next day, the cells were co-transfected with pSin-DsRed-IRES-Puro (or the GFP-expressing equivalent), the psPAX2 packaging vector, and one of the Spike-expressing PB plasmids depending on the experiment performed. A 1:1:1 molar ratio of all three plasmids for a total DNA concentration of 20 μg or 40 μg (T75 vs. T150 flask) was used to transfect the cells using the TransIT^®^-LT1 Transfection Reagent (Mirus Bio). Three days later, the supernatant was collected and spun at 350xg for 10 minutes at 4° C, then filtered through a 0.45 μM filter. The virus was then pelleted by ultracentrifugation at 100,000 x g over a 20% sucrose cushion, as described previously ^19^. The virus was then quantified using the Lenti-X™ p24 Rapid Titer Kit (Takara Bio) and aliquots were frozen at −80° C for future use.

### Inducible cell line production

HEK293T cells were plated at ~50% confluence in a 6-well plate one day before transfection. The cells were transfected the next day with pTransposase and PB-Tet-hAce2 at a molar ratio of 1:4. 2.5 μg of total DNA was used for the transfection. Two days later, the cells were split into a 10-cm plate, and Blasticidin was added to select stably transfected cells at a concentration of 10 μg/ml. The selected cells were grown in the presence of Blasticidin for one more week. The cells were then induced with 1000 ng/ml of doxycycline for 48 hours. To select for the inducible cell colonies that express ACE2 at a higher level, we then stained with 2 μg of anti-ACE2 goat IgG antibody (R&D Systems, AF933) per million cells followed by a FITC-conjugated secondary antibody (Santa Cruz Biotechnology, sc-2356). The unbound antibodies were washed away. The positive gate for sorting was chosen to eliminate the cells expressing low levels of ACE2 based on low FITC signal. The sorted cells were grown out in the presence of Blasticidin until ready for experimentation.

### In vitro infection assay

30,000 of the HEK-ACE2 cells were plated in each well of a flat-bottom 96-well plate. The indicated concentration of virus was added to each well containing 100 μl of media (DMEM +10% FBS) in triplicates. After overnight incubation at 37° C, the media was removed from the cells without disturbing the adherent cell layer. 200 μl of fresh media was added back to the cells. The cells were then incubated in 37°C for an additional 24 hours. The cells were resuspended by pipetting and washed in wash media (PBS + 2% BSA) once and resuspended in the same media. Flow cytometry was used to analyze the infection.

For the in vitro competition assay, we added 5.0 ng p24 equivalent of the RFP-expressing S^D614^-pseudotyped virus to each well of a flat-bottom 96-well plate containing 30,000 HEK-ACE2 cells. At the same time, we added increasing concentrations of either the GFP-expressing S^D614^- or S^G614^-pseudotyped viruses, each in triplicates. The plates were then placed at 4° C to allow equilibration for one hour, followed by incubation at 37° C. On day one post-infection, we removed the supernatant and replaced it with fresh media. 24 hours later, we quantified the infection by each of the GFP-expressing and RFP-expressing viruses using flow cytometry.

For the infection of the ACE2-expressing inducible cell lines, the sorted cells were plated in 6-well plates at a concentration of 500,000 cells per plate. The cells were treated with indicated concentrations of doxycycline for two days. Subsequently, the cells were removed with 5% EDTA solution and replated at a density of 30,000 cells per well in a flat-bottom 96-well plate while maintaining the same doxycycline concentration as in the 6-well plate. 7.5 ng or 15 ng of RFP-expressing S^D614^- or S^G614^-pseudotyped virus was added to the cells at each level of induction. The infection was done in triplicates. The cells were incubated at 37° C for 48 hours, and the RFP was quantified using the flow cytometer as before.

### Virus crosslinking and Spike trimer purification

We adopted the protocol previously described for crosslinking and purification of the HIV-1 glycoprotein trimers ^20^ to the SARS-CoV-2 Spike trimers. Briefly, 1740 ng p24 equivalent of each virus type was resuspended in PBS in a total volume of 100 μl. Bis(sulfosuccinimidyl) (BS3, ThermoFisher Scientific, 21580) at a final concentration of 2mM was added to crosslink the Spike trimers. After 20 minutes at room temperature, 20 mM of Tris-HCl (pH 7.5) was added and mixed by pipetting to quench the crosslinking reaction. The virus was pelleted by centrifugation (20,000 x g for 45 minutes), and the supernatant was aspirated and discarded. The pelleted virus was resuspending by pipetting in 100 μl of PBS. A mild detergent, n-Dodecyl β-D-maltoside (DDM, D4641, Sigma), at a final concentration of 0.1% was used to lyse the virus.

Magnetic beads with covalently bound anti-FLAG (Sigma, M8823) or anti-HA (ThermoFisher Scientific, 88838) antibodies were used for immunoprecipitation of the crosslinked trimers using their C-terminal tags. The beads were washed once in IP wash media (0.1% DDM in PBS). The crosslinked and lysed virus samples were divided into two equal volumes, and 10 μl of each bead type was added to each of the samples followed by incubation at room temperature for 3 hours with continuous rotation. The beads were washed using 1 ml of the wash media by rotation at room temperature for 5 minutes, followed by placing them adjacent to a magnet for 2 minutes and removing the supernatant containing the unbound proteins. The process was repeated three more times. The bound trimers were eluted using 50 μl of the elution buffer provided in the HA immunoprecipitation kit (ThermoFisher Scientific, 88838), which is a low pH buffer (pH = 2.0). After 10 minutes of incubation, 7.5 μl of the neutralization buffer, also provided in the same kit, was added. The samples were then run on a gel, and western blot analysis was performed as described below.

### Western blotting and protein quantification

Cell lysates were prepared using Radioimmunoprecipitation assay (RIPA) lysis buffer supplemented with protease inhibitors (cOmplete Protease inhibitor cocktail, Sigma-Aldrich, 11697498001) and 50mM 1,4-Dithiothreitol (DTT). Protein extracts were quantified using Bio-Rad DC Protein Assay kit. For western blot analysis, 50 μg of protein was mixed with NuPAGE 4x loading buffer, heated to 95° C for 10 minutes, and then run on NuPAGE 4-12% polyacrylamide gels (Invitrogen) and transferred to a Polyvinylidene difluoride (PVDF) membrane using Bio-Rad Trans-Blot^®^ SD semi-dry transfer blotting apparatus. For viral protein analysis, 355 ng of p24 equivalent of each virus was used. Blots were blocked for 10 minutes using 5% non-fat dry milk in Tris-buffered saline (TBS) containing 0.05% Tween. Blots were incubated with primary antibodies for either one hour at room temp or overnight at 4°C with rocking. HRP conjugated secondary antibodies were used at 1/5000 dilution for 30 minutes. The blots were developed using Western Lightning Plus ECL reagent from Perkin Elmer.

### Antibodies

In addition to the antibodies named in the previous sections, the following antibodies were used for western blot analysis: Rabbit anti-GAPDH (2118) and rabbit anti-HA (3724) were purchased from Cell Signaling and used at 1/1000 dilution. Mouse anti-HIV-1 integrase (SC-69721) was purchased from Santa Cruz Biotechnology and used at 1/5000 dilution. Mouse anti-Flag antibody M2 was purchased from Sigma and used at 1/1000. HRP conjugated secondary antibodies were purchased from Pierce.

### Statistical analysis

A two-tailed Student’s t-Test was used to ascertain statistical differences. The built-in software in Microsoft^®^ Excel^®^ was used.

## Results

### The D614G mutation increases the entry fitness of Spike pseudotyped lentiviral particles

We developed a lentiviral vector pseudotyped with the Spike protein of the SARS-CoV-2 to evaluate the entry fitness of the different Spike variants. The Spike protein’s C-terminal domain contains a signal sequence that targets the nascent protein to the intermediate compartment between the endoplasmic reticulum and the Golgi complex (ERGIC) ^15,21^. However, efficient pseudotyping of the lentiviral vectors requires glycoproteins’ presence on the cell membrane at the time of viral budding. To find the Spike modification that leads to the most efficient pseudovirus production, we tested three different strategies: unmodified Spike protein, the 19-terminal amino acids deleted, and finally, the terminal 19-amino acids of the Spike protein replaced with the HIV-1 gp41 signal sequence (NRVRQGYS) ^15^ (Supplementary Figure 1A). In the virus-producing cells, the expression level of the unmodified Spike protein and the Spike protein with the gp41 signal peptide was the highest. However, in the pseudotyped particles, the version with the 19 terminal amino acids deleted had higher Spike protein levels (Supplementary Figure 1B and 1C). We then compared the infectivity of the lentiviral vectors pseudotyped with either of the three Spike protein modifications (Supplementary Figure 1D). In addition, we tested the ACE2-dependence of each of these three viruses by comparing the infection of the HEK293T cells to the ACE2-expressing HEK-ACE2 cells (Supplementary Figure 1E). Based on these results, for the remainder of this work we chose the Spike protein version with the 19 C-terminal amino acids deleted.

Single-round lentiviral particles pseudotyped with either S^D614^ or S^G614^ were made and quantified as described in the methods section. From this point on, the S^D614^-pseudotyped lentiviral vectors will be referred to as D614 virus and the S^G614^-pseudotyped virus will be referred to as G614 virus. The pseudotyped viruses were created using a GFP-expressing backbone plasmid, Psin-GFP-IRES-Puro. We used increasing concentrations of the pseudotyped lentiviral vectors to infect HEK-ACE2 cells. These cells are a monoclonal cell line generated by exogenous transduction of ace2 into HEK293T cells and selection for high expression levels of ACE2. ^14,15^. Two days after the infection, we quantified the percent of infected cells by flow cytometry. Increasing the concentration of the viruses pseudotyped with either of the Spike variants increased the percentage of infected cells; however, compared to the D614 virus, the G614 virus infected more cells for a given virus concentration. At the maximum virus concentration used (50 ng p24 equivalent), both the D614 and G614 viruses lead to high levels of infection and the difference between them was only marginally significant (61.3% vs. 68.4%, respectively, *p*-value = .06) (Figure 1A). However, at lower concentrations, the G614 virus infectivity was significantly higher than the D614 virus (Figure 1A). At 6.2 ng p24 equivalent of the virus, the G614 virus infectivity was 3.6-fold higher than that of the D614 virus (Figure 1B).

**Figure 1:**
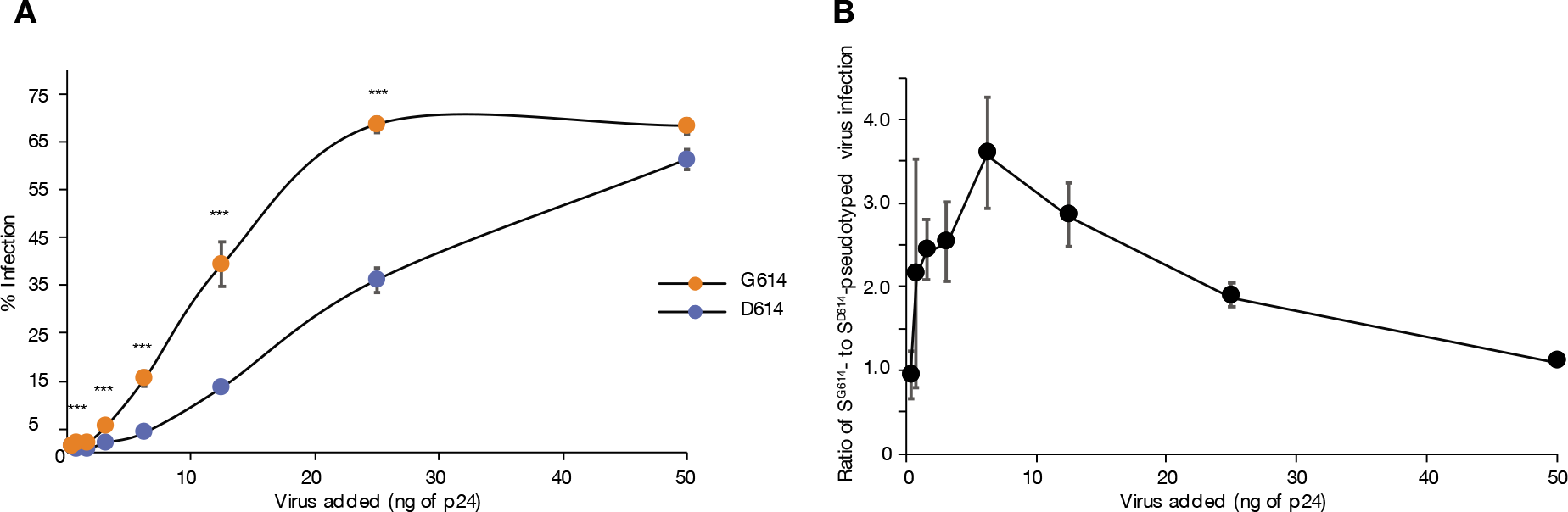
Entry fitness of G614 compared to the D614 variant. **A)** Percent of HEK-ACE2 cells infected with S^D614^- or S^G614^-pseudotyped lentivirus, as a function of amount of virus added (ng of p24). Both pseudoviruses expressed GFP, and the percentage of GFP-positive cells was quantified by flow cytometry two days after infection. The infection was done in triplicates. **B)** The ratio of the percentage of the cells infected by the G614 virus to the percentage of the cells infected by the same amount of the D614 variant. *, **, and *** indicate p-values ≤ .05, .01, and .001, respectively.

### The G614 variant is capable of outcompeting the D614 variant in an in vitro competition assay

While the previous experiment and other similar reports ^1,7^ suggest that the entry of the G614 virus is up to 3.6-fold higher than that of the D614 virus, the physiological consequence of this increased entry is unclear. In particular, it is unclear if this mutation can outcompete the original D614 variant if both viruses infect the same host at the same time. Answering this question would require animal models of the disease or more granular viral dynamics data in patients. We attempted to address this question by an in vitro competition assay in which viral variants compete to infect target cells, and the G614 virus can bind to and replace the D614 virus from the surface of the ACE2-expressing cells.

In this assay, cells were co-incubated with a fixed concentration of an RFP-expressing D614 virus and an increasing concentration of the GFP-expressing G614 virus. The plates were then placed at 4° C to allow equilibration for one hour, followed by incubation at 37°C for two days (Figure 2A). As a control experiment, increasing concentrations of a GFP-expressing D614 virus was added to the cells along with a fixed concentration of the same variant (D614) expressing RFP. In both experiments, we measured the fraction of all infected cells that were RFP^+^ (i.e. infected with RFP-D614 virus only) as a function of the amount of GFP-expressing virus added (D614 or G614) (Figure 2B). As expected, we observed that for higher levels of GFP-expressing virus added, the resulting fraction of infected cells expressing RFP was lower. More importantly, we found that at every input ratio of GFP:RFP virus, the G614-GFP virus was more efficient at outcompeting the D614G-RFP than the control D614G-GFP variant was. RFP^+^ and GFP^+^ infected cells occurred at approximately equal frequencies if a ~1:1 inoculum of D614-RFP and D614-GFP virus were added, however, if the same inoculum ratio was done with G614-GFP, then <20% of infected cells were RFP+. The G614-GFP virus could replace ~50% of D614-containing RFP^+^ infected cells even when it was inoculated at only ~1:3 ratio. These results suggest that when encountering the same ACE2-expressing cell, the G614 variant is able to directly and efficiently outcompete the D614 virus.

**Figure 2:**
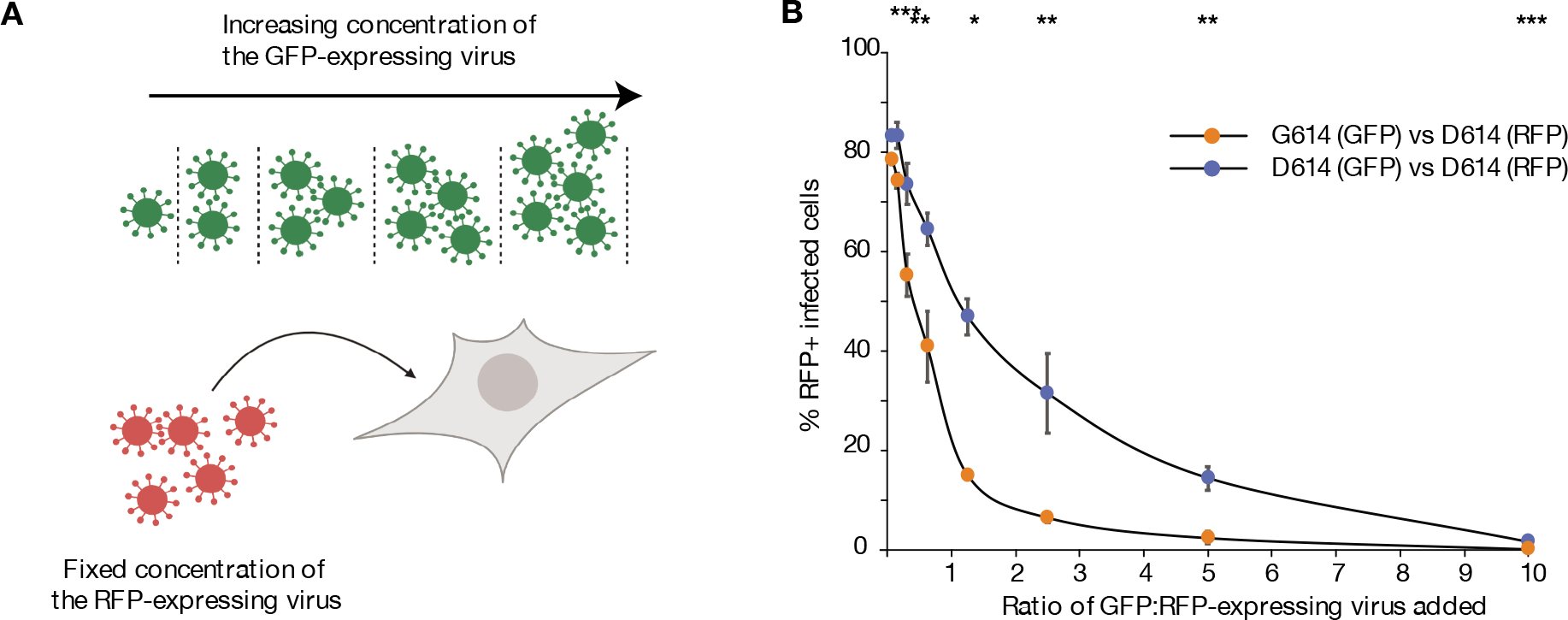
In vitro competition assay. **A)** Schematic of experimental design. A fixed concentration of the RFP-expressing D614 virus was added to the HEKACE2. At the same time, increasing concentrations of the GFP-expressing D614 or G614 virus was added to the cells to outcompete the RFP-expressing D614 virus. After 1 hour of incubation at 4° C to allow equilibration, the cells were incubated at 37 degrees for 2 days. The percentage of the cells expressing GFP and RFP was quantified using flow cytometry. **B)** Outcomes of the competition assay. At each ratio of GFP-expressing to RFP-expressing virus added, the percent of total infected cells (RFP^+^ or GFP^+^) that were RFP^+^ is reported. Lower percentage of RFP^+^ cells imply that the GFP-expressing virus was more efficient at binding to and replacing the RFP-expressing virus from the surface of the HEKACE2 cells. The competition between GFP-expressing G614 and D614 virus is shown in orange. The control competition of GFP-expressing D614 with RFP-expressing D614 virus is shown in blue. *, **, and *** indicate p-values ≤ .05, .01, and .001, respectively.

### The G614 entry fitness advantage over the D614 variant is independent of the level of ACE2 expression

Sequencing analysis of human-derived tissues has identified a variety of primary cells that express the ACE2 receptors and other co-factors necessary for the infection ^12,22^. Ex vivo infectivity assays have confirmed the infectibility of some of these cells by SARS-CoV-2^23–25^. The surface expression level of ACE2 on these cells varies greatly^26^. Moreover, given the central role of ACE2 in the Renin-Angiotensin system, its expression varies on each cell type over time to maintain the overall fluid balance and blood pressure ^27^. Therefore, depending on the cell type being infected and the homeostatic state of the body, the Spike protein mediates entry into cells with widely different ACE2 expression levels. In contrast, the entry fitness advantage of the G614 compared to the D614 variant has so far only been studied in cell lines artificially induced to express high levels of exogenous ACE2 or non-human primate cell lines expressing the ACE2 receptor ^7,14,15^.

To investigate the fitness advantage of the D614G Spike mutation under different ACE2 expression levels, we made an inducible cell line expressing ACE2 under the control of doxycycline. We used an all-in-one, Tet-On 3G inducible PiggyBac (PB) plasmid, as described in the Methods section. We co-transfected HEK293T cells with the inducible plasmid and a second plasmid encoding the PB transposase enzyme. The transposase mediates the integration of the ace2-containing gene cassette into the cellular genome. The cassette also contains the blasticidin resistance gene, and thus, the stably transfected cells can be selected with blasticidin. After one week of selection, the un-transfected cells died out. Inducible cells were then treated with 2 μg of doxycycline for 48 hours and stained for ACE2 expression. The selected cells were first stained with an anti-ACE2 antibody, followed by a GFP-expressing secondary antibody. The cells with the highest expression of the ACE2, as judged by the most intense GFP signal, were sorted. The sorted cells were then expanded in culture in the absence of doxycycline for an additional week.

Next, we established a dose-response-curve for the doxycycline-mediated induction of the ACE2 expression on the sorted cells. Cells were treated with doxycycline, and the percentage of ACE2-expressing cells was determined using surface staining and flow cytometry (Figure 3A). At 80 ng/ml of doxycycline, 33.9% of the inducible cells were ACE2 positive. Higher concentrations of doxycycline did not significantly increase the ACE2 expression level and negatively influenced their viability. Meanwhile, only 1.49% of the cells treated with Dimethyl Sulfoxide (DMSO) were positive for ACE2 expression. In parallel, the HEK293T cells obtained from Michael Farzan ^14^ showed 64.5% positivity for ACE2 expression by flow cytometry (Figure 3B). In addition, the surface expression level of the ACE2, as measured by mean fluorescent intensity (MFI) when stained with an anti-ACE2 antibody, also increased with increasing doxycycline concentrations (Figure 3C).

**Figure 3:**
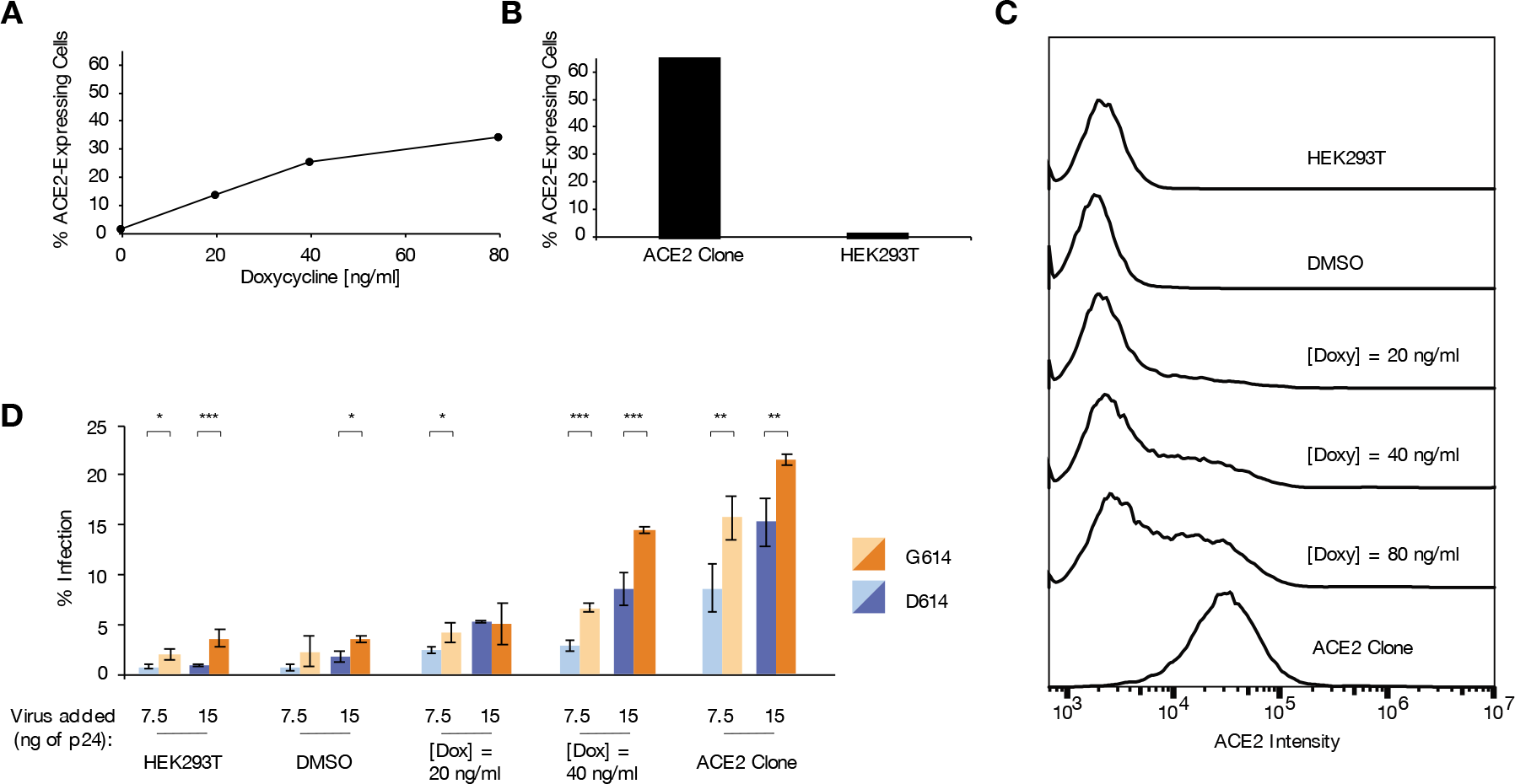
The entry fitness advantage of the G614 relative to the D614 variant is independent of the amount of ACE2 expressed. An HEK293T cell line expressing ACE2 under the control of a doxycycline-inducible promotor was created and used for infection assays. **A)** The percentage of the inducible cells expressing ACE2 as a function of the concentration of doxycycline added, quantified using surface staining and flow cytometry. **B)** The percentage of cells expressing ACE2 in the the parental HEK293T cells and the cell line previously engineered for high ACE2 expression (HEK-ACE2). Cell surface staining and flow cytometry analysis was done similar to panel B and is described in the Methods section. **C)** The distribution of ACE2 expression levels on the surface of the inducible cells, parental HEK293T cells, and the HEK-ACE2 cells. Cells were stained with an anti-ACE2 antibody followed by a GFP-expressing secondary antibody as in panel B and C. The mean of fluorescent intensity (MFI) in the GFP channel was used to quantify the ACE2 expression level. **D)** Percent of cells infected with Spike-pseudotyped virus for different Spike variants (G614 vs D614), viral inoculum sizes (7.5 vs 15 ng), and cellular ACE2 expression levels. Infection assays used 30,000 cells and were done in triplicate. Infection was quantified by RFP expression 2 days after virus was added, using flow cytometry. *, **, and *** indicate p-values ≤ .05, .01, and .001, respectively.

We then infected inducible cells expressing a range of surface ACE2 levels with either D614 or G614 viruses. The infection was done in triplicates using two different concentrations of each virus (7.5 ng and 15 ng of p24 equivalent per 30,000 cells in a flat-bottom 96-well plate). Both the D614 and G614 viruses infected a greater number of cells as the concentration of doxycycline used to induce the ACE2 expression increased (Figure 3D). Moreover, independent of the ACE2 expression levels, the G614 virus infected more cells than the D614 virus (Figure 3D). For example, at 40 ng/ml of doxycycline, 25% of the inducible cells express ACE2 on their surface. At this level of ACE2 expression, 7.5 ng p24 of the G614 virus infects the cells 2.3 times more efficiently than the D614 virus (6.7% vs. 2.86%, p-value = 0.0005). Interestingly, even in uninduced cells with very low ACE2 expression levels similar to that of the parental HEK293T cells (1.49% vs. 1.34%, respectively), there was some infection of the cells with virus pseudotyped by both Spike variants and the infection level was significantly higher for the G614 variant compared to the D614 variant. In this case, 7.5 ng p24 of the G614 variant infects 2.3% of the un-induced cells, compared to 0.61% infection achieved with the same concentration of the D614 variant (p-value 0.019). Even the parental HEK293T not expressing any exogenous ACE2 can be infected with S-pseudotyped lentiviral vectors: 1.98% and 0.75% with the G614 and D614 variants, respectively (Figure 3D).

### Total and surface expression of the G614 and D614 Spike variants are equal in virus-producing cells, but viral incorporation of the G614 Spike protein is greater

The previous experiments suggest that the G614 variant has a higher entry fitness than the D614 variant in our in vitro pseudotyping assay and that the fitness advantage is independent of the level of ACE2 expression on the surface of the cells. The D614G mutation occurs 66 amino acids upstream of the polybasic S1/S2 Furin cleavage site and 109 amino acids downstream of the receptor-binding domain (RBD) of the S1 subunit ^3^. Since the D614G mutation is not in the vicinity of the RBD of Spike, it is unlikely that the observed increase in fitness occurs by a direct change in the interaction between the Spike trimers and the ACE2 receptor. In preprint work, Zhang and colleagues ^7^ have demonstrated an increase in the number of Spike monomers and S1 subunit on the surface of the virions pseudotyped with G614 compared to the D614 variant. This finding points to an increase in the avidity rather than the affinity of the G614 Spike trimers for the ACE2 receptor. We sought first to confirm this finding in our system and second to determine why more G614 Spike proteins are packaged in the virions compared to the D614 variants. One possibility is more efficient production or surface expression of the Spike proteins on the surface of the viral-producing cells.

A higher overall expression of the S^G614^ compared to the S^D614^ could lead to increased overall viral incorporation and may explain the higher entry fitness. We transfected HEK293T cells with a plasmid expressing either the D614 or the G614 variant of the S protein. 48 hours later, we lysed the cells using RIPA buffer and performed western blot analysis under denaturing conditions. We used an anti-HA antibody to detect the C-terminal tagged Spike protein and an anti-Glyceraldehyde 3-phosphate dehydrogenase (GAPDH) antibody for total protein loading control. We performed the transfection and western blot analysis in three independent replicates (Figure 4A). We quantified the expression level of the 70-kDa S2 subunit and the 180-kDa un-cleaved S monomer (Figure 4B). The un-cleaved S expression was slightly lower in the G614 variant than the D614 variant (adjusted band intensity to GAPDH (ABI) 0.95 vs. 1.05, p-value = .009), but the amount of the cleaved S2 fragment was not significantly different between the two variants (ABI 1.09 vs. 0.91, p-value = .36).

**Figure 4:**
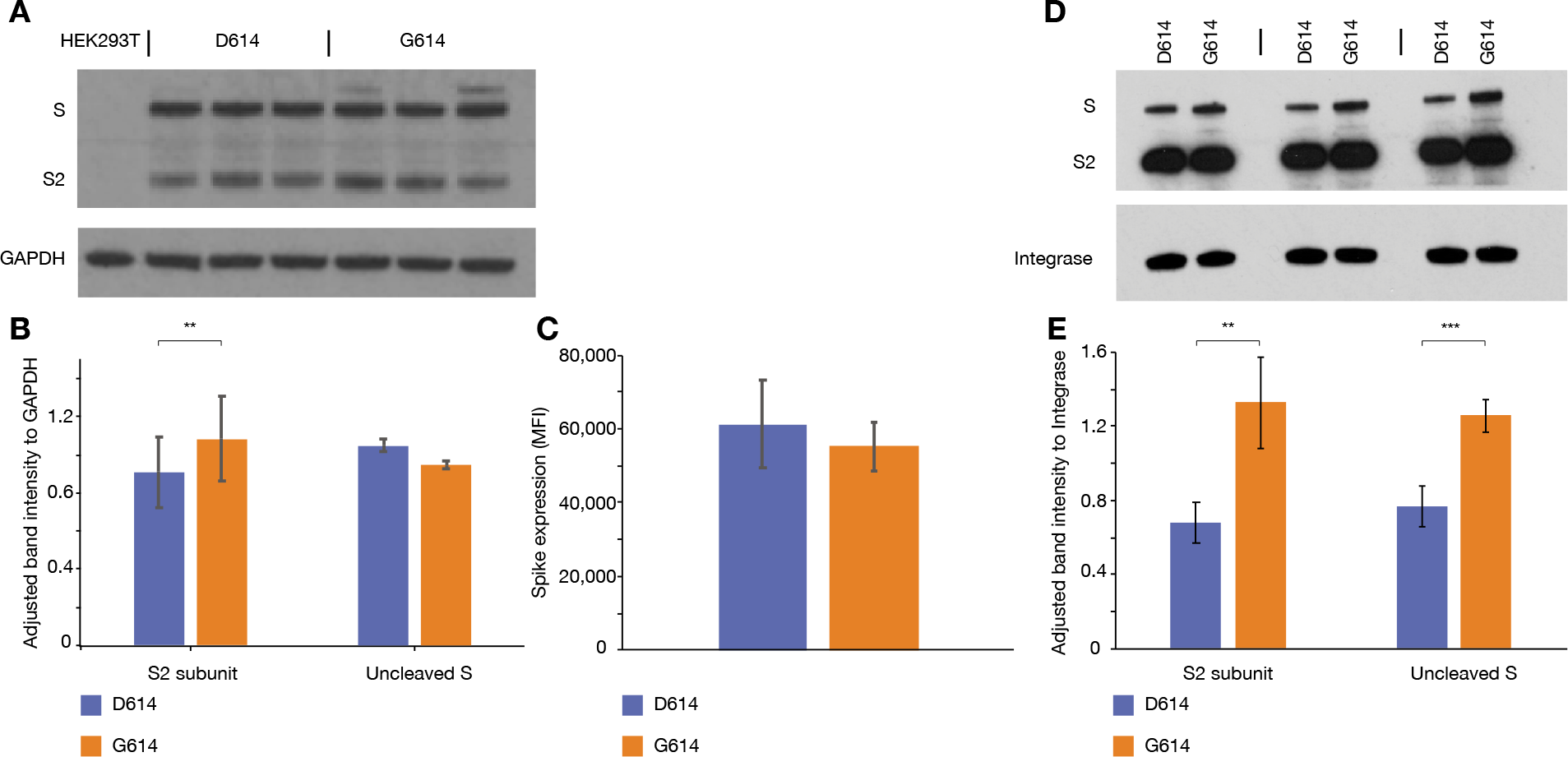
Total and surface expression of the different Spike variants in virus-producing cells as well as their incorporation into the virions. **A)** Expression of Spike variants in virus-producing cells. HEK293T cells were transfected with plasmids encoding the D614 and G614 variants of the Spike protein with a fused C-terminal HA tag. Western blot analysis with anti-HA antibody was done to quantify the amount of un-cleaved S and cleaved S2 subunit present in the cell lysate. The transfection and western blot analysis were done in triplicates. **B)** The amount of the S and S2 subunits was quantified relative to the signal intensity of the GAPDH band. **C)** The surface expression of each Spike variant on the virus-producing cells. Cells were transfected with plasmids encoding the different Spike variants, D614 and G614 as before. After 2 days, the transfected cells were incubated with a solubilized ACE2 molecule carrying a human IgG1 Fc at the C-terminus, followed by a PE-conjugated mouse-anti-human secondary antibody. The amount of each Spike variant protein on the transfected cells’ surface was quantified as the MFI in the PE channel. The transfection and staining was done in triplicates. **D, E)** The amount of each Spike variant incorporated into the virion. D614 and G614 viruses were made and concentrated as before. 355 ng p24 equivalent of each virus was lysed and loaded on the gel for western blot analysis using anti-HA antibody. **E)** The amount of un-cleaved S and cleaved S2 subunit was quantified relative to the Integrase band’s signal intensity. Three independent replicates were analyzed. *, **, and *** indicate p-values ≤ .05, .01, and .001, respectively.

We next examined if the S protein’s surface expression on the virus-producing cells is higher for the G614 compared to the D614 variant of the spike protein. Such a difference could also lead to higher incorporation of the Spike protein in the G614 to the D614 viruses and a higher entry fitness. We first transfected HEK293T cells with plasmids expressing the S^G614^ and S^D614^ proteins. 48 hours later, we stained the cells with a solubilized ACE2 molecule carrying an IgG1 Fc at the C-terminus, followed by a secondary mouse anti-human antibody conjugated to R-phycoerythrin (PE). The Spike protein expression level was quantified as the mean of the fluorescent intensity in the PE channel (MFI) using flow cytometry (Figure 4C). There was no significant difference between the surface expression of S^G614^ and S^D614^ (MFI 55,198 vs. 61,232. p-value = .57). Therefore, neither the total protein production nor the surface expression of the Spike variants is significantly different.

Finally, we examined if the Spike protein incorporation into virions is different between the S^G614^ and S^D614^ variants. D614- or G614 viruses were made as described before. Three days after transfection, the virus was concentrated by ultracentrifugation over 20% sucrose cushion. We first verified that this method of viral concentration excludes free proteins in the supernatant that could influence analysis of the proteins incorporated into the virions. We collected samples at three different stages of the ultracentrifugation step: 10 μl of the supernatant prior to the spin, 10 μl from the top fraction after ultracentrifugation, and 10 μl of the virus pellet resuspended in 200 μl of PBS after decanting the rest of the supernatant. Both the pre-spin supernatant and the top fraction after the ultracentrifugation contained free Integrase (both fully cleaved and partially-cleaved Pol molecules) and Spike proteins. However, the viral pellet obtained after ultracentrifugation over a 20% sucrose cushion only contains fully processed proteins present in the virus particle (Supplemental Figure 2). Next, we proceeded to quantify each of the Spike variants present in the viral pellet. 355 ng p24 equivalent of the virus was lysed and loaded for WB analysis under denaturing conditions. We used an anti-HA antibody for quantification of the Spike protein and anti-Integrase antibody for ensuring equal loading of each viral preparation. Western blots were done in triplicates (Figure 4D). We quantified the relative band intensity of the cleaved S2 or un-cleaved S proteins adjusted to the Integrase intensity (Figure 4E). Consistent with prior unreviewed reports ^7^, compared to the D614 virus, the G614 virus had a higher amount of both the cleaved S2 subunit (adjusted band intensity relative to the Integrase (ABI) of 1.33 vs. 0.68, p-value = .01) as well as the un-cleaved S polyprotein (ABI of 1.26 vs. 0.77, p-value = .004).

### Chimeric Spike trimers containing both types of Spike protomers are enriched in the G614 compared to the D614 variant

CryoEM analysis suggests that the D614 and T859 residues from neighboring spike monomers form a hydrogen bond ^1,4,28^, and that the D614G mutation abrogates this bond. The change in the non-covalent bonds between the neighboring monomers could fundamentally change the Spike trimers’ stability. It has been shown that the Spike proteins on the surface of coronaviruses are almost exclusively in the form of trimers composed of individual Spike protomers^29^. However, the prior work quantifying the different Spike variants incorporated in the virions was done under denaturing conditions that destabilize the non-covalent bonds holding the Spike trimers together. While such protocols can elucidate the total number of Spike monomers in the virions, the more physiologically relevant question concerning the number and makeup of the Spike trimers remains unaddressed.

We hypothesized that the higher concentration of the S^G614^ compared to the S^D614^ monomers in the virions could be the result of the greater propensity of the G614 monomer to form stable trimers and get incorporated into the virion. To test this, we synthesized chimeric virions packaging both the D614 and G614 Spike variants and quantified the amount of each monomer and trimer. To distinguish between the S^D614^ and S^G614^ proteins packaged in the chimeric virions, we used different C-terminal tags, 3xFLAG and HA, respectively. As a control, chimeric virions expressing the FLAG-tagged S^D614^ and HA-tagged S^D614^ were also synthesized. To make these viruses containing chimeric Spike trimers, HEK293T cells were transfected with plasmids expressing both Spike versions at a 1:1 molar ratio along with the other two plasmids necessary to make virus. The resulting viruses were collected and concentrated as before. Viruses were lysed, and the amount of each Spike variant was analyzed with western blot using anti-HA and anti-Flag antibodies normalized to the amount of Integrase (Figure 5A). As shown in Figure 5B, there were twice as many HA-tagged Spike proteins in the D614-FLAG/G614-HA chimeric virions than the D614-FLAG/D614-HA virions (1.5 vs. 0.7, p-value =.06). On the other hand, there was no significant difference in the amount of FLAG-tagged Spike protein between these two chimeric virus types (Figure 5C). Therefore, in the situation where two different viral strains infect the same cell, despite the equal expression level of D614 and G614 variants (Figure 4C), the G614 variant is preferentially packaged in the budding virion.

**Figure 5:**
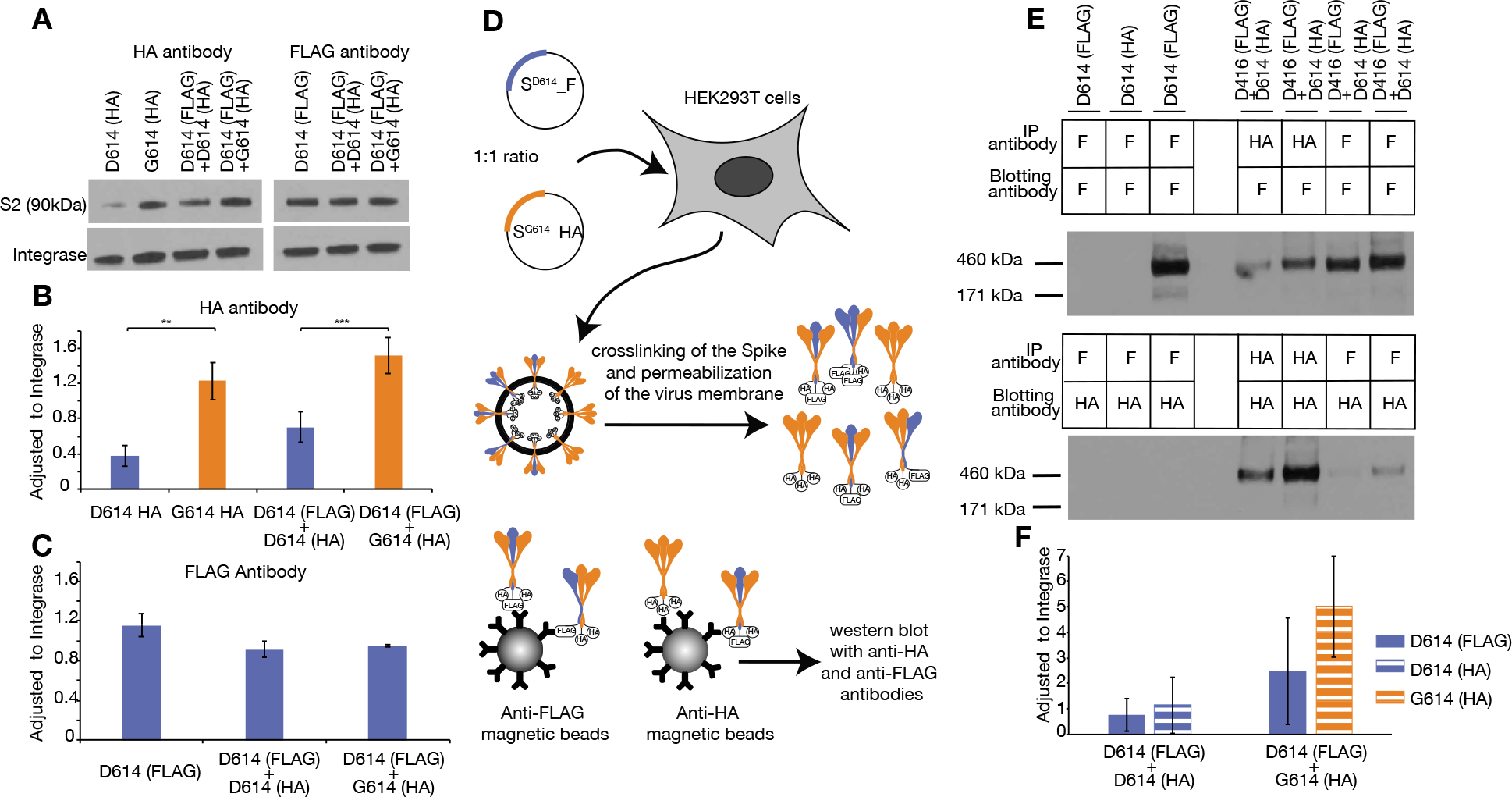
The differential trimerization of the D614 vs. G614 Spike monomers. **A, B, C)** Differential packaging of Spike monomers into viruses produced from cells containing equal amounts of S^D614^ and S^G614^. Equal molar ratios of plasmids making D614-FLAG and G614-HA or D614-FLAG and G614-HA were transfected into HEK293T cells along with the other necessary constructs to make virions with a mix of Spike proteins. The resulting viruses were collected as before and lysed for western blot analysis. **B, C)** The western blot analysis for the FLAG-tagged Spike proteins and the HA-tagged Spike proteins was done in triplicates and normalized to the Integrase content. **D)** Schematic of the protocol used to stabilize and purify the Spike trimers from the surface of the virions. The viruses containing both the FLAG-tagged D614 Spike and either the HA-tagged G614 or the HA-tagged D614 were made from HEK293T cells as described above. The trimers were crosslinked by treating the viral prep with BS3 followed by permeabilization with DDM. The crosslinked Spike trimers were purified by IP using either anti-HA- or anti-FLAG-conjugated magnetic beads. **E)** Different amounts of each Spike variant are present in chimeric Spike trimers. The vertically-oriented text above the table denotes the virus from which the Spike trimer is purified. The magnetic bead used to purify the Spike trimer is indicated in the first column: F: anti-FLAG-conjugated beads, HA: anti-HA-conjugated beads. The purified trimers were then analyzed by western blot using anti-Flag (top blot) or anti-HA (bottom blot) antibody. **F)** The relative amount of G614 vs. D614 monomers present in chimeric mutant/wild-type trimers. Anti-Flag-conjugated magnetic beads were used to purify the extracted chimeric Spike trimers. The amount of D614 monomer was quantified (solid blue bar). In addition, the amount of HA-tagged D614 (blue striped bar) and HA-tagged G614 (orange striped bar) was also quantified from Spike trimers extracted from the virions and purified with anti-FLAG beads. The IP and WB analysis were done in triplicates. The Spike protein amounts were normalized to the amount of intra-virion Integrase. D614(HA): viruses containing HA-tagged D614 Spike. D614(FLAG): virus containing the 3xFLAG-tagged Spike. D614 (FLAG)+D614(HA): viruses containing both the 3xFLAG-tagged Spike and the HA-tagged D614 Spike. D614 (FLAG)+G614(HA): viruses containing both the 3xFLAG-tagged D614 Spike and the HA-tagged G614 Spike. *, **, and *** indicate p-values ≤ .05, .01, and .001, respectively.

Next, we confirmed that the trimeric Spike proteins on the surface of the virions made when both versions of the Spike protein are present is indeed a chimera containing both monomers. This requires stabilization of the Spike trimers. To do so, we used a previously-described protocol for the HIV-1 glycoprotein trimer ^20^ to stabilize the trimers. Chimeric virions were made containing both the S^D614^-FLAG and S^G614^-HA. As a control, we also made virions pseudotyped with both S^D614^-FLAG and S^D614^-HA. The trimers were crosslinked using bis(sulfosuccinimidyl)suberate (BS3). BS3 is an amine-to-amine crosslinker and has been shown to maintain the three-dimensional structure of the viral glycoproteins and maintain their antigenicity^20^. The virus was then lysed with n-Dodecyl-B-D-Maltoside (DMM), a maltoside-based non-ionic detergent. Next, we separated the HA- or FLAG-tagged Spike trimers from the other viral proteins by immunoprecipitation (IP) with magnetic beads covalently bound to anti-HA or anti-FLAG antibodies (Figure 5B). After elution of the bound Spike trimers, we analyzed the quantity of the different Spike variants. We could purify trimers from the D614-FLAG/G614-HA chimeric virions with either anti-FLAG or anti-HA magnetic beads. Western blot analysis showed that these virions contained both HA- and FLAG-tagged Spike proteins (Figure 5E), proving that chimeric Spike trimers containing both S^D614^ and S^G614^ were formed. In contrast, trimers isolated from virions pseudotyped with only the D614-FLAG, D614-HA, or G614-HA Spike proteins could only be purified and blotted with the appropriate antibodies corresponding to the C-terminal tag of their respective Spike.

Next, using chimeric viruses containing both Spike variants, we sought to compare the propensity of the S^G614^ compared to the S^D614^ to form Spike trimers. FLAG- and HA-based immunoprecipitation could have different efficiencies due to the difference in the anti-FLAG and anti-HA antibodies’ affinities. To account for these intrinsic differences in the IP efficiency, we quantified the Spike variants in chimeric trimers pulled down with anti-FLAG beads. The D614-FLAG/G614-HA chimeric trimers contained 4.4 times more HA-tagged Spike proteins, representing the G614 variant, compared to the D614-FLAG/D614-HA trimers pulled down with the same beads (Figure 5F, chemiluminescence signal intensity of 5.0 vs. 1.1 after adjusting to the virus input using Integrase band intensity). Therefore, in a system where both the D614 and G614 Spike variants are synthesized and expressed with equal efficiencies, the trimerized Spike proteins packaged in budding virions contain more of the G614 variant compared to the D614 variant. This preferential trimerization could explain the previous observation that the viruses pseudotyped with S^G614^ contain more Spike protein than the S^D614^-pseudotyped viruses. More spike trimers on the surface of the viruses increases the avidity for the ACE2 receptor on target cells and can lead to the greater entry fitness of the G614 variant.

## Discussion

The D614G mutation in the SARS-CoV-2 Spike protein characterizes a distinct viral lineage that emerged during the 2020 pandemic and become the dominant variant globally. The gradual replacement of the D614 by the G614 variant and the higher viral load of the patients infected with the virus carrying the D614G mutation suggests that the mutation confers fitness advantage on the virus ^1^. In this study we investigated the functional significance of this mutation in a cell culture system using lentiviruses expressing different Spike variants on their surface. We found that the D614G Spike mutation has a fitness advantage over the original variant for infecting ACE2-expressing cells, in agreement with published work by Korber et al.^1^ and a pre-print by Zhang et al^7^. We also showed that in a direct competition assay, the G614 variant is superior to the D614 variant in terms of competing for ACE2 receptors and entering target cells. Furthermore, using an inducible cell line with ACE2 expression levels under the control of doxycycline, we showed that the fitness advantage of the G614 compared to the D614 variant is independent of the amount of ACE2 expressed on the surface of the cells. Therefore, as the SARS-CoV-2 target different cells of the body, each with different surface expression levels of ACE2, the fitness advantage of the G614 over the D614 variant is likely maintained. We observed a trend towards increased Spike proteins on the surface of the G614 virus vs the D614 virus, agreeing with Zhang et al. ‘s work ^7^. Finally, we showed that compared to D614, the G614 variant is preferentially present in chimeric trimers formed when a mixture of both variants is present in the virus-producing cells. Therefore, despite the loss of the hydrogen bond between the D614 and T859 on the neighboring Spike protomers, the G614 variant is likely to produce more stable trimers.

One limitation of our findings is the use of in vitro infectivity assays. Although we sought to improve the system’s physiological relevance by utilizing cells with variable and controllable expression levels of ACE2, our results are limited to pseudotyped lentiviruses and thus may differ from physiological settings. Furthermore, similar to other groups ^7,9^, in order to increase the efficiency of the Spike protein expression on the surface of the virus-producing cells, and therefore, the overall infectivity, we modified the Spike protein by deleting the C-terminal 19 amino acids and instead inserted HA or FLAG peptide tags. This modification modifies the cognate signal peptide on the Spike and preferentially targets the nascent Spike proteins to the cell’s surface []. Therefore, the experiments looking at the Spike protein’s packaging in the virions could be affected by this artificial trafficking. However, we synthesized both the D614 and G614 Spike variants with similar C-terminus modifications, and therefore, the comparison between the amount of these two proteins may still reflect their relative amount in the unmodified SARS-CoV-2. Finally, while our system can be used to investigate the relative amount of each Spike variant in the chimeric trimers, further work is needed to determine if the different Spike trimers have different three-dimensional configurations.

In addition to characterizing the fitness advantage of the D614G mutation of the Spike protein and the mechanism by which this advantage is obtained, we have developed several techniques that can be used more broadly by the research community. The ACE2 expression level varies widely depending on the cell type being infected and the homeostatic condition of the body. Despite this, most prior studies utilize cells with fixed, and often supraphysiologic levels of ACE2 expression ^14,15^. To our knowledge, we have established the first cell line with an inducible level of ACE2 expression that can be used to study the dependence of the SARS-CoV-2 on ACE2. Furthermore, developing and testing neutralizing antibodies against the Spike trimer requires the stabilization of the trimers. Other groups have achieved this by introducing mutations or exogenous trimerization peptides in the Spike protein sequence ^30–32^. Such mutations may result in changes in the three-dimensional structure of the Spike trimers and alter their antigenicity. Here we have adopted a chemical crosslinking technique borrowed from the HIV-1 field ^20^ for stabilization and purification of Spike proteins expressed on the surface of pseudoviruses. This technique does not alter the Spike protein sequence and has been shown to maintain the antigenicity of the HIV-1 glycoprotein trimers ^20^.

## Acknowledgements

We are extremely grateful to the expert review of Dr. Alison Hill and her gracious help with editing of this work. We also would like to acknowledge Dr. Sahand Hormoz for his invaluable feedback on the experimental methods.

## Author Contributions

WAM designed and performed the experiments and assisted in preparation of the manuscript. GMB designed the experiments and assisted in preparation of the manuscript.

SAR conceived the study, planned and performed the experiments and wrote the manuscript.

## Reference

1. Korber B, Fischer WM, Gnanakaran S, et al. Tracking changes in SARS-CoV-2 Spike: evidence that D614G increases infectivity of the COVID-19 virus. Cell. Published online July 2020:S0092867420308205. doi:10.1016/j.cell.2020.06.043

2. Hoffmann M. SARS-CoV-2 Cell Entry Depends on ACE2 and TMPRSS2 and Is Blocked by a Clinically Proven Protease Inhibitor. :19.

3. Andersen KG, Rambaut A, Lipkin WI, Holmes EC, Garry RF. The proximal origin of SARS-CoV-2. Nat Med. Published online March 17, 2020. doi:10.1038/s41591-020-0820-9

4. Yuan M, Wu NC, Zhu X, et al. A highly conserved cryptic epitope in the receptor-binding domains of SARS-CoV-2 and SARS-CoV. Science. Published online April 3, 2020. doi:10.1126/science.abb7269

5. Song W, Gui M, Wang X, Xiang Y. Cryo-EM structure of the SARS coronavirus spike glycoprotein in complex with its host cell receptor ACE2. PLoS Pathog. 2018;14(8). doi:10.1371/journal.ppat.1007236

6. Farkas C, Fuentes-Villalobos F, Garrido JL, Haigh J, Barría MI. Insights on early mutational events in SARS-CoV-2 virus reveal founder effects across geographical regions. PeerJ. 2020;8:e9255. doi:10.7717/peerj.9255

7. Zhang L, Jackson CB, Mou H, et al. The D614G Mutation in the SARS-CoV-2 Spike Protein Reduces S1 Shedding and Increases Infectivity. Microbiology; 2020. doi:10.1101/2020.06.12.148726

8. Yurkovetskiy L, Pascal KE, Tompkins-Tinch C, et al. SARS-CoV-2 Spike protein variant D614G increases infectivity and retains sensitivity to antibodies that target the receptor binding domain. bioRxiv. Published online July 4, 2020. doi:10.1101/2020.07.04.187757

9. Daniloski Z, Guo X, Sanjana NE. The D614G mutation in SARS-CoV-2 Spike increases transduction of multiple human cell types. bioRxiv. Published online June 15, 2020. doi:10.1101/2020.06.14.151357

10. Ogawa J, Zhu W, Tonnu N, et al. The D614G mutation in the SARS-CoV2 Spike protein increases infectivity in an ACE2 receptor dependent manner. bioRxiv. Published online July 22, 2020. doi:10.1101/2020.07.21.214932

11. Chow RD, Chen S. The Aging Transcriptome and Cellular Landscape of the Human Lung in Relation to SARS-CoV-2. Genomics; 2020. doi:10.1101/2020.04.07.030684

12. Zhao Y, Zhao Z, Wang Y, Zhou Y, Ma Y, Zuo W. Single-cell RNA expression profiling of ACE2, the putative receptor of Wuhan 2019-nCov. bioRxiv. Published online January 26, 2020:2020.01.26.919985. doi:10.1101/2020.01.26.919985

13. Sungnak W, Huang N, Bécavin C, et al. SARS-CoV-2 entry factors are highly expressed in nasal epithelial cells together with innate immune genes. Nature Medicine. Published online April 23, 2020:1–7. doi:10.1038/s41591-020-0868-6

14. Li W, Moore MJ, Vasilieva N, et al. Angiotensin-converting enzyme 2 is a functional receptor for the SARS coronavirus. Nature. 2003;426(6965):450–454. doi:10.1038/nature02145

15. Moore MJ, Dorfman T, Li W, et al. Retroviruses Pseudotyped with the Severe Acute Respiratory Syndrome Coronavirus Spike Protein Efficiently Infect Cells Expressing Angiotensin-Converting Enzyme 2. Journal of Virology. 2004;78(19):10628–10635. doi:10.1128/JVI.78.19.10628-10635.2004

16. Yu J, Vodyanik MA, Smuga-Otto K, et al. Induced pluripotent stem cell lines derived from human somatic cells. Science. 2007;318(5858):1917–1920. doi:10.1126/science.1151526

17. Frieda KL, Linton JM, Hormoz S, et al. Synthetic recording and in situ readout of lineage information in single cells. Nature. 2017;541(7635):107–111. doi:10.1038/nature20777

18. Randolph LN, Bao X, Zhou C, Lian X. An all-in-one, Tet-On 3G inducible PiggyBac system for human pluripotent stem cells and derivatives. Sci Rep. 2017;7. doi:10.1038/s41598-017-01684-6

19. Rabi SA, Laird GM, Durand CM, et al. Multi-step inhibition explains HIV-1 protease inhibitor pharmacodynamics and resistance. J Clin Invest. 2013;123(9):3848–3860. doi:10.1172/JCI67399

20. Leaman DP, Lee JH, Ward AB, Zwick MB. Immunogenic Display of Purified Chemically Cross-Linked HIV-1 Spikes. Journal of Virology. 2015;89(13):6725–6745. doi:10.1128/JVI.03738-14

21. Sadasivan J, Singh M, Sarma JD. Cytoplasmic tail of coronavirus spike protein has intracellular targeting signals. J Biosci. 2017;42(2):231–244. doi:10.1007/s12038-017-9676-7

22. Qi F, Qian S, Zhang S, Zhang Z. Single cell RNA sequencing of 13 human tissues identify cell types and receptors of human coronaviruses. Biochemical and Biophysical Research Communications. 2020;526(1):135–140. doi:10.1016/j.bbrc.2020.03.044

23. Kim HK, Kim H, Lee MK, et al. Generation of tonsil organoids as an ex vivo model for SARS-CoV-2 infection. bioRxiv. Published online August 7, 2020:2020.08.06.239574. doi:10.1101/2020.08.06.239574

24. Yang L, Han Y, Nilsson-Payant BE, et al. A Human Pluripotent Stem Cell-based Platform to Study SARS-CoV-2 Tropism and Model Virus Infection in Human Cells and Organoids. Cell Stem Cell. 2020;27(1):125–136.e7. doi:10.1016/j.stem.2020.06.015

25. Zhang B-Z, Chu H, Han S, et al. SARS-CoV-2 infects human neural progenitor cells and brain organoids. Cell Research. Published online August 4, 2020:1–4. doi:10.1038/s41422-020-0390-x

26. Ziegler CGK, Allon SJ, Nyquist SK, et al. SARS-CoV-2 Receptor ACE2 Is an Interferon-Stimulated Gene in Human Airway Epithelial Cells and Is Detected in Specific Cell Subsets across Tissues. Cell. 2020;181(5):1016–1035.e19. doi:10.1016/j.cell.2020.04.035

27. Vaduganathan M, Vardeny O, Michel T, McMurray JJV, Pfeffer MA, Solomon SD. Renin–Angiotensin–Aldosterone System Inhibitors in Patients with Covid-19. N Engl J Med. Published online March 30, 2020:NEJMsr2005760. doi:10.1056/NEJMsr2005760

28. Walls AC, Park Y-J, Tortorici MA, Wall A, McGuire AT, Veesler D. Structure, Function, and Antigenicity of the SARS-CoV-2 Spike Glycoprotein. Cell. Published online March 9, 2020. doi:10.1016/j.cell.2020.02.058

29. Delmas B, Laude H. Assembly of coronavirus spike protein into trimers and its role in epitope expression. Journal of Virology. 1990;64(11):5367–5375. doi:10.1128/JVI.64.11.5367-5375.1990

30. Juraszek J, Rutten L, Blokland S, et al. Stabilizing the Closed SARS-CoV-2 Spike Trimer. bioRxiv. Published online July 10, 2020:2020.07.10.197814. doi:10.1101/2020.07.10.197814

31. McCallum M, Walls AC, Bowen JE, Corti D, Veesler D. Structure-guided covalent stabilization of coronavirus spike glycoprotein trimers in the closed conformation. Nature Structural & Molecular Biology. Published online August 4, 2020:1–8. doi:10.1038/s41594-020-0483-8

32. Xiong X, Qu K, Ciazynska KA, et al. A thermostable, closed SARS-CoV-2 spike protein trimer. Nature Structural & Molecular Biology. Published online July 31, 2020:1–8. doi:10.1038/s41594-020-0478-5

