## Supplemental Table and Figures for "The SARS-CoV-2 Spike mutation D614G increases entry fitness across a range of ACE2 levels, directly outcompetes the wild type, and is preferentially incorporated into trimers"

### Supplementary Material

| Plasmid | Sequence (5’-> 3’) |
|---|---|
| CoV-10 | ggcctttcgacctgcagcccaagcttatgtttgtgttcctg |
| CoV-20 | gcctttcgacctgcagcccaagcttatggagctgaggccctgg |
| CoV-24 | tcacgcataatccggcacatcatacggataacaacaggagccacaggaac |
| CoV-21 | ccgatttaaattcgaattcgctagcttagagggcgctctggtc |
| CoV-31 | ctaagaacctgaatgagtccc |
| CoV-35 | ctcagtacagttcacaccctggtagagcacagc |
| CoV-36 | gctgtgctctaccagggtgtgaactgtactgag |
| CoV-37 | ggagattctggacatcacacc |
| CoV-39 | ggtaccatgtcaagctcttcctggctccttc |
| CoV-40 | actagtctaaaaggaggtctgaacatcatcag |
| CoV-42 | atgatctttataatcaccgtcatggtctttgtagtcacaacaggagccacaggaac |
| CoV-43 | tcacttgtcatcgtcatccttgtaatcgatgtcatgatctttataatcacc |
| CoV-45 | tgcagcccaagcttatgtttgtgttcctggtg |
| CoV-46 | gtacctagctagctcacttgtcatcgtcatcc |

**Table 1: Sequence of the primers and oligos used.**

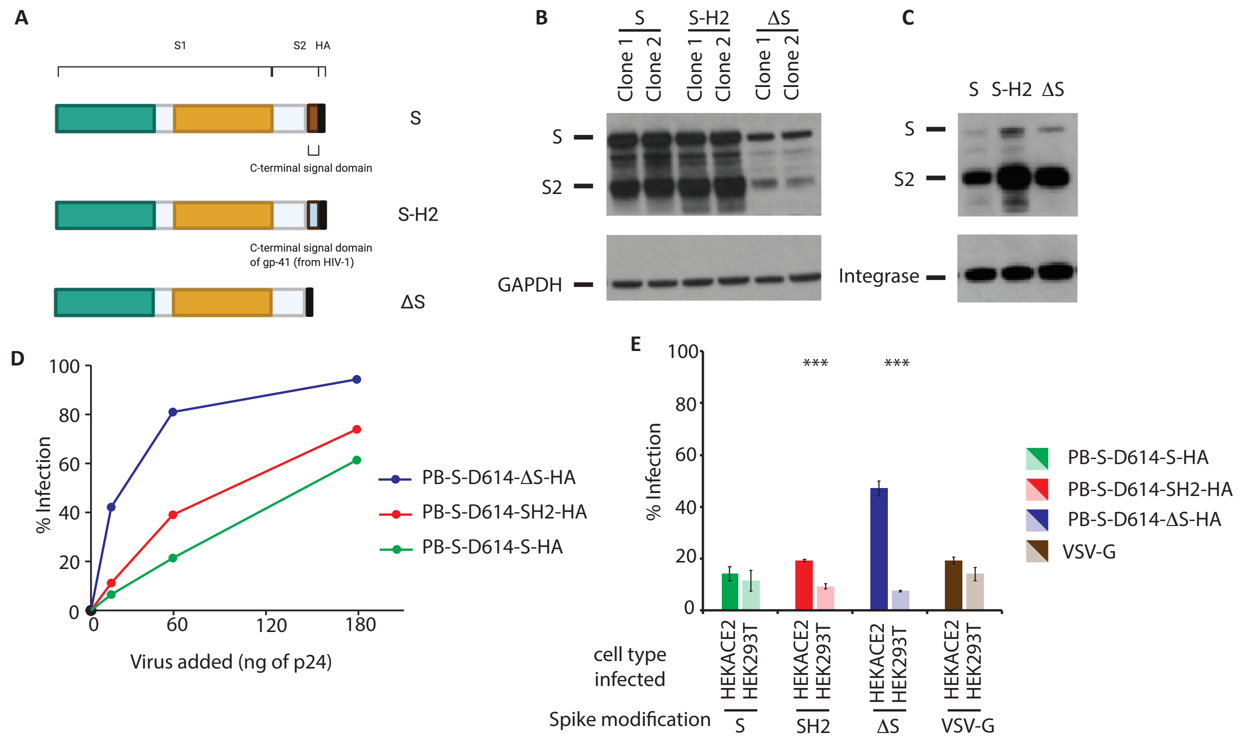


**Supplementary Figure 1**: Choice of the Spike modification for lentiviral-based pseudotyping. **A)** Schematic of the different Spike modifications tested. S: C-terminal signal sequence not modified, S-H2: The C-terminal signal sequence was deleted and replaced with the signal sequence of the HIV-1 gp41. ΔS: The C-terminal signal sequence was deleted. All three Spike protein modifications were tagged with an HA peptide. **B)** Expression of the different Spike protein modifications in virus-producing cells. After transfection with plasmids expressing the different Spike modifications, S, S-H2, and ΔS, the cells were lysed, and the total amount of the Spike protein was detected using an anti-HA antibody. GAPDH was used for loading control. **C)** Incorporation of the Spike proteins with different modifications within lentiviruses. Three different pseudoviral preparations were made as described in the Methods Section. Each was made using a different Spike modification: S, S-H2, and ΔS. The viruses were lysed, and the total amount of the Spike protein was detected using an anti-HA antibody. Intra-virion Integrase was used for loading control. **D)** Infectivity of the lentiviruses pseudotyped with different Spike modifications. The three different lentiviruses described above were used to infect HEK-ACE2 cells. The viruses express RFP, and the percentage of the cells infected is measured for each virus using flow cytometry. **E)** ACE2 dependence of the lentiviruses pseudotyped with different Spike modifications. Either HEK293T cells or the HEK-ACE2 cells were infected with the above lentiviruses. As before, the percentage of each cell type infected with these viruses was measured using flow cytometry. The infections were done in triplicates. *** indicates p-value < .001.

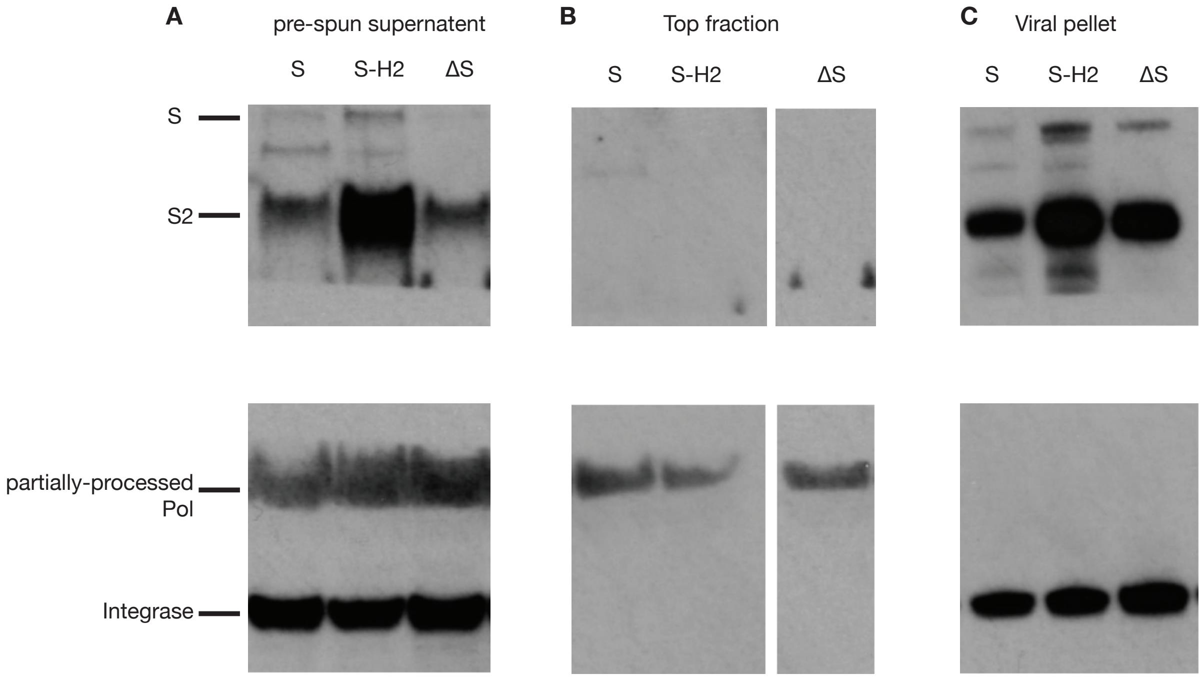


**Figure 2.** Analysis of the virion-associated and free Spike and Integrase proteins. HEK293T cells were transfected to make Spike-pseudotyped lentiviruses as described in the Methods section. The C-terminal of the Spike protein is tagged with an HA peptide. **A)** Three days after transfection 10 µl of the supernatant was collected (pre-spun supernatant) and the amount of Spike proteins (un-cleaved S and cleaved S2) and fully-processed Integrase enzyme (32kDa) or its intermediate cleavage product (partially-processed Pol at ~ 55 kDa) was analyzed using western blot analysis. The rest of the supernatant was ultracentrifuged over a 20% sucrose cushion, as described previously. After ultracentrifugation, 10 µl of media was pipetted from the top. **B)** The presence of the same proteins was analyzed in this fraction. Next, the supernatant was completely decanted, and the virus pellet was resuspended in 100 µl of PBS. **C)** 10 µl of the virus was loaded and analyzed by western blot for the same proteins. Note: in panel B, the ΔS lane was loaded on a different gel and transferred to a different blot. The image was cropped, and the lane was placed next to the other two lanes. In addition, panel C is the identical image to panel C of the Supplementary Figure 1 and is placed here for direct comparison with the other two viral prep fractions. The three different lanes represent lentiviruses pseudotyped without modification of the Spike C-terminal signal sequence (S), Spike with the 19 C-terminal amino acids representing the signal peptide deleted and replaced with the signal peptide of the HIV-1 gp-41 (S-H2) or the 19 C-terminal amino acids deleted (ΔS).
